# The ClpP Peptidase Forcefully Grips Protein Substrates

**DOI:** 10.1101/2022.05.14.491966

**Authors:** Steven D. Walker, Adrian O. Olivares

## Abstract

ATPases Associated with diverse cellular Activities (AAA+) proteases power the maintenance of protein homeostasis by coupling ATP hydrolysis to mechanical protein unfolding, translocation, and ultimately degradation. Though ATPase activity drives a large portion of the mechanical work these molecular machines perform, how the peptidase contributes to the forceful denaturation and processive threading of substrates remains unknown. Here, using single-molecule optical trapping, we examine the mechanical activity of the Caseinolytic Peptidase P (ClpP) from *Escherichia coli* in the absence of a partner ATPase and in the presence of an activating small molecule acyldepsipeptide. We demonstrate that ClpP grips protein substrate under mechanical loads exceeding 40 pN, which are greater than those observed for the AAA+ unfoldase ClpX and the AAA+ protease complexes ClpXP and ClpAP. We further characterize substrate-ClpP bond lifetimes and rupture forces under varying loads. We find that the resulting slip bond behavior does not depend on ClpP peptidase activity. Additionally, we find that unloaded bond lifetimes between ClpP and protein substrate are on a timescale relevant to unfolding times (up to ∼160 s) for difficult to unfold model substrate proteins. These direct measurements of the substrate-peptidase bond under load define key properties required by AAA+ proteases to mechanically unfold and degrade protein substrates.

**STATEMENT OF SIGNIFICANCE:** Energy-dependent proteases drive essential protein degradation to maintain cellular homeostasis and to rapidly regulate protein levels in response to changes in cellular environment. Using single-molecule optical tweezers, several studies demonstrate that the molecular process of degradation involves the mechanical unfolding and translocation of protein substrates by the ATP hydrolyzing enzyme component of these protease complexes. This study provides evidence that the chambered peptidase component of these molecular machines also contributes to the mechanical process of degradation by gripping substrate under load in a manner independent of peptide hydrolysis. Our results suggest that the peptidase actively contributes to the biophysical mechanisms underlying processive protein degradation by energy-dependent proteolytic machines.

## INTRODUCTION

AAA+ (<u>A</u>TPases <u>A</u>ssociated with diverse cellular <u>A</u>ctivities) proteases power protein degradation in the cell to eliminate damaged or misfolded proteins and control cellular processes by modulating protein levels. These proteolytic molecular machines comprise ATP-dependent, ring-shaped motor proteins (i.e. AAA+ protein unfoldases) that recognize, unfold and translocate substrate protein into a self-compartmentalized peptidase (1, 2). In *Escherichia coli*, the ClpP peptidase pairs with the AAA+ unfoldases ClpA and ClpX to form functional AAA+ proteases (3, 4). Importantly, protein degradation is processive (i.e., enzyme translocates along protein substrate without dissociating until degradation is complete). Robust processivity requires that the probability of substrate dissociation is low and that subunits within the translocating machinery maintain enzymatic cycles out of phase such that polypeptide does not dissociate during substrate unfolding and translocation.

Single-molecule studies in the past decade have illuminated how the AAA+ proteases ClpXP and ClpAP function during processive protein unfolding and translocation. More specifically, optical trapping experiments combined with solution biochemistry revealed how AAA+ proteases generate force, coordinate ATPase cycles, grip protein substrate, and translocate along the polypeptide track (5–13). Because the AAA+ unfoldase behaves as a motor protein (i.e., couples chemical energy in the form of ATP hydrolysis to physical translocation along a macromolecular track) and unfolds and translocates substrate protein in the absence of its proteolytic partner, the peptidase has been overlooked as contributing to the chemomechanical cycle of protein degradation by AAA+ proteases. In fact, there is little direct evidence that ClpP generates force during the process of protein degradation. However, the AAA+ motors show mechanical defects when ClpP is not present. For example, ClpA unfolds a dimeric substrate more slowly (14) and takes slower kinetic steps in the absence of ClpP (15, 16). Moreover, ClpX, which grips substrate via its pore-1 loops (10, 11, 17), slips on substrate more often than ClpXP (10). Though differences in motor function can be partially explained by changes in ATPase activity in the presence of ClpP (18, 19), we sought to test if ClpP participates mechanically during protein degradation since direct observation of ClpP mechanics has not been demonstrated.

Obtaining direct evidence of ClpP mechanical degradation has been complicated as ClpP poorly degrades large unfolded protein substrates in the absence of AAA+ motors due to gating by its N-terminal loops (20, 21). AAA+ unfoldase binding activates ClpP by opening the axial pore and allowing polypeptide to be threaded into the ClpP chamber for degradation. However, a class of natural products called acyldepsipeptides (ADEPs) activate ClpP in the absence of motor proteins (22, 23), allowing the direct observation of protein degradation by ClpP (Fig. 1 A). ADEPs bind to the same hydrophobic pocket on ClpP that motors bind (24) and are thought to activate ClpP in a similar manner, i.e. by causing rearrangement of the ClpP N-terminal loops (25). In cells, this activation leads to indiscriminate proteolysis and ultimately cell death, making ClpP a promising target for developing antibiotics that hamper pathogenic biofilm formation (26) and anti-cancer therapies due to ClpP’s conserved role in mitochondrial protein homeostasis (27, 28). Additionally, ADEPs disrupt bacterial cell division through the specific degradation of the protein FtsZ by ClpP (29). Furthermore, using purified proteins, ADEP-activated ClpP appears to unfold and degrade FtsZ *in vitro* without the need for motor proteins (30). Taken together, these data suggest that ClpP is capable of mechanically engaging and degrading substrate in the absence of a AAA+ unfoldase.

**FIGURE 1.**
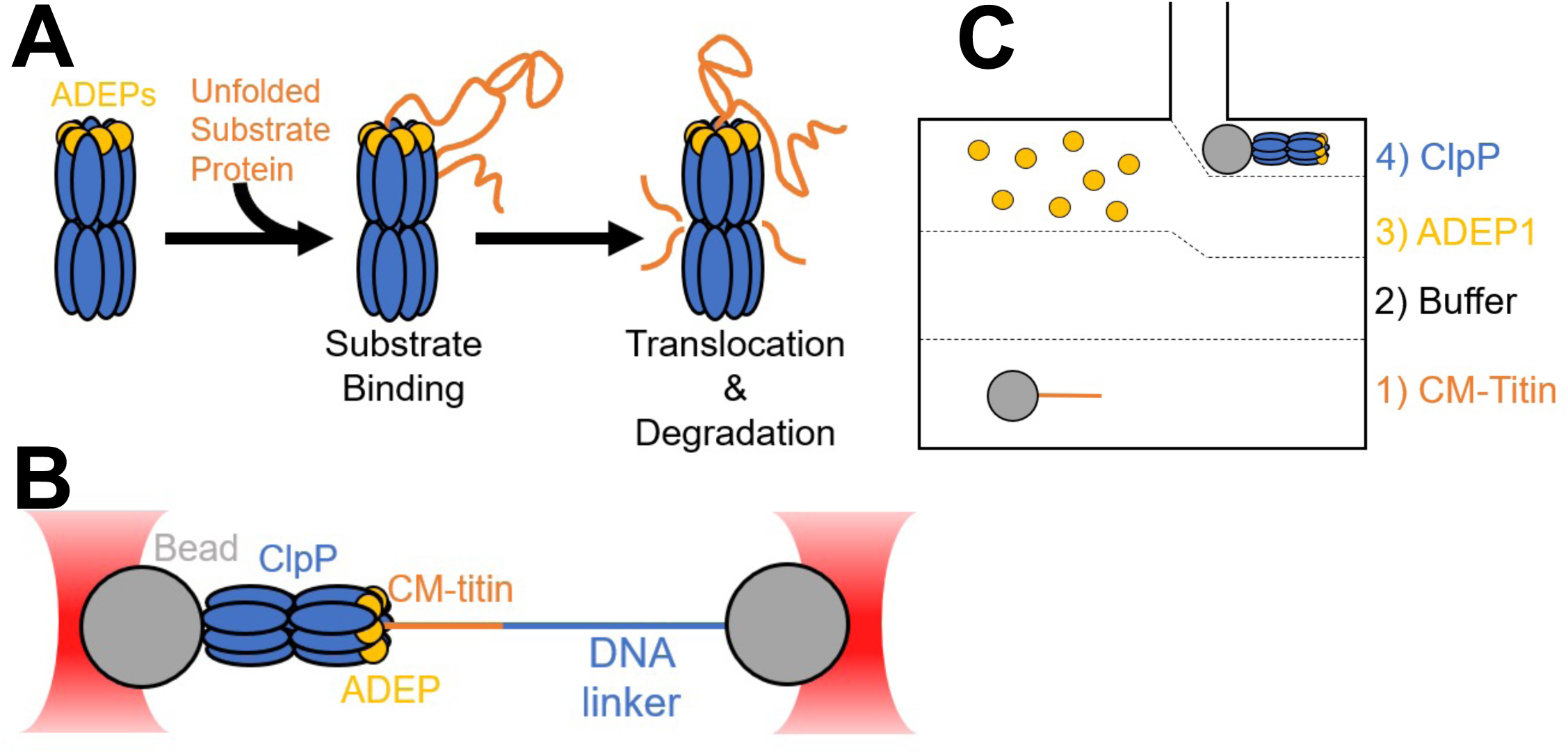
Measuring single-molecule ClpP mechanics by optical trapping. *(A)* Cartoon of ADEP activation of ClpP. ADEPs (yellow) bind to the ClpP tetradecamer (blue), opening its central pore and allowing metastable protein substrates (orange) to enter and be degraded. *(B)* Schematic of optical trapping assays. ClpP is immobilized to beads and engages a CM-titin substrate bound beads using a 3500bp DNA linker. *(C)* Example schematic of the flow cell used for optical trapping showing a top-down view. Solutions were prepared separately and flown into the channels as shown. Single substrate- and ClpP-coated beads were captured in channels 1 and 4, respectively. The stage was then moved to channel 3 where experiments were performed in the presence of 10 µM ADEP.

Here, we aim to provide evidence of ClpP’s contribution to mechanical protein degradation in the absence of motor proteins. We hypothesize that ClpP grips and degrades substrate against external force, and that the active site serine contributes to substrate grip. Using single-molecule optical trapping, we do not observe denatured substrate translocation but demonstrate that ClpP grips substrate against applied loads in excess of 40 pN when activated by ADEP. We find that substrate-ClpP bond lifetimes and rupture forces decrease as external load increases consistent with slip bond behavior. We further show that active site inactivation does not significantly affect substrate grip and discuss what other portions of ClpP likely account for its mechanical behavior. To our knowledge, this study provides the first direct evidence that ClpP maintains a force-dependent grip on protein substrates without a motor protein and suggests additional activities that ClpP may contribute to ATP-dependent processive translocation and protein degradation outside of its peptidase activity.

## MATERIALS AND METHODS

### Biochemical purification of ClpP and substrate proteins

Full length *E*. coli ClpP with a C-terminal hexahistidine tag and a terminal cysteine residue, and a substrate protein comprising an N-terminal HaloTag domain, four variant titin^I27^ domains (V13P), and a C-terminal hexahistidine tag and 11-amino acid ssrA degron were cloned and purified as previously described (31). Briefly ClpP was cloned into the pQE70 plasmid and expressed in JK-10 cells, which lack endogenous ClpP (32). Cells were initially grown to OD_600_ ∼0.6 in LB broth at 30°C, cooled to 18°C, and induced with 0.5 mM IPTG for expression overnight. Cells were harvested and resuspended in lysis buffer (50 mM sodium phosphate pH 8.0, 1 M NaCl, 5 mM imidazole, 10% glycerol), frozen in liquid nitrogen, and stored at −80°C. For purification, all the following steps were performed at 4°C unless noted otherwise. Cells were thawed and lysed with two passes through an Emulsiflex high-pressure homogenizer (Avestin, Canada). The lysate was clarified by centrifugation at 30,000 x g for 30 minutes. Clarified lysate was passed through an INDIGO-Ni (Cube Biotech, Germany) affinity column, washed, and eluted with lysis buffer containing 500 mM imidazole. Fractions were analyzed by SDS-PAGE, pooled, and concentrated to ∼1 mL. Concentrated protein was further purified by size exclusion chromatography using a HiPrep 16/60 Sephacryl S300-HR column (Cytiva) equilibrated with storage buffer (50 mM Tris-HCl pH 8.0, 150 mM KCl, 0.5 mM EDTA, 10% glycerol). Fractions were analyzed by 12% SDS-PAGE and appropriate fractions pooled, concentrated using an Amicon Ultra-15 10kDa MWCO centrifugal filter (Millipore Sigma), flash frozen and stored at −80°C. ClpP was biotinylated at the terminal C-terminal cysteine using EZ-Link^™^ Maleimide-PEG2-Biotin (Thermo Scientific). First a 20 mM stock of the biotin-maleimide was made in storage buffer and added to a final concentration of 20x molar excess to ClpP. The sample was left rotating at 4°C overnight and buffer exchanged into storage buffer containing 1 mM DTT before freezing with liquid nitrogen. Protein concentration was determined in storage buffer using ε_280_ =125,160 M^-1^ cm^-1^ for the ClpP tetradecamer.

The substrate protein was cloned into a pFN18A plasmid (Promega) and expressed in BL21(DE3) cells. Cells were grown to OD_600_ ∼0.6 in LB broth at 37°C, cooled to 25°C, and induced with 1 mM IPTG for 3 hours. Cells were harvested by centrifugation at 4000 x g for 15 min, resuspended in lysis buffer (50 mM Sodium phosphate pH 8.0, 500 mM NaCl, 10% glycerol, 10 mM β-mercaptoethanol, 20 mM imidazole) and flash frozen in liquid nitrogen for storage at −80°C. Lysis, clarification, and INDIGO-Ni affinity was performed as described above eluting with 250 mM imidazole. Fractions were analyzed by 12% SDS-PAGE. Pure fractions were pooled, concentrated, flash frozen in small aliquots. Protein concentration was determined using ε_280_ =89,380 M^-1^ cm^-1^. For titin domain carboxymethylation, aliquots of substrate were first unfolded using 2 M guanidine-HCl at room temperature for 1.5 hours. Then, a fresh stock of 0.5 M iodoacetic acid was added to a final concentration of 2.5 mM. After another 1.5-hour incubation, the reaction was quenched by adding excess 1 M DTT to a final concentration of 10 mM. Samples were buffer exchanged into PD buffer (25 mM Hepes pH 7.6, 100 mM KCl, 10% glycerol, 10 mM MgCl_2_, 0.1% Tween-20) and flash frozen for storage at −80°C.

### Single-molecule optical trapping of ClpP-substrate complexes

For trapping experiments, biotinylated ClpP and substrate were immobilized onto 1.25-micron streptavidin beads (Spherotech) in PD buffer supplemented with 1 mg/mL BSA (PD-BSA). For substrate, we constructed a 3500 bp linker with a 3’ 20 bp overhang from the M13mp18 plasmid (Bayou Biolabs) by PCR using these primers (Integrated DNA Technologies): TTTCCCGTGTCCCTCTCGA-T/idSp/TTGAAATACCGACCGTGTGA, and AATCCGCTTTGCTTCTGAC with a 5’ biotin. The complement to the 20 bp overhang with sequence ATCGAGAGGGACACGGGAAA contained a 5’ phosphate and 3’ amine to which a HaloTag substrate was conjugated using a HaloTag succinimidyl ester O4 ligand (Promega). The 3500 bp DNA linker was ligated in the presence of CM-titin at room temperature for >1 hour before conjugating to streptavidin beads.

All optical trapping data were collected using a dual-laser m-Trap Optical Tweezers system (LUMICKS) equipped with a 5-channel laminar flow microfluidics device. Prior to experiments, the microfluidic chip was washed extensively with ddH_2_ O and equilibrated with PD-BSA for >30 min. ClpP and substrate beads were washed and resuspended in PD-BSA containing an oxygen scavenging system (0.25 mg/mL glucose oxidase, 0.03 mg/mL catalase, 3 mg/mL glucose; PD-BSA-OX). ClpP-bound beads and substrate beads were flown into the 2^nd^ and 5^th^ channels, with the 3^rd^ and 4^th^ containing PD-BSA-OX and PD-BSA-OX supplemented with 10 μM ADEP1 (Cayman Chemical), respectively. Custom Python scripts were written and used to automate bead capture, force ramp/clamp control, and to perform data analysis. For rupture force experiments, after capturing beads with each trap and forming a tether, the script moved the beads at a constant velocity until a rupture occurs. Similarly, for lifetime traces the script steered the trap until a defined force is reached, after which it paused until the tether ruptures. Traces with incorrect contour lengths or multiple ruptures were discarded to avoid non-specific interactions and multiple tethers.

### Optical trapping data analysis

Data analysis for both lifetime and rupture force experiments were carried out using custom Python scripts with the LUMICKS Bluelake software. For rupture force measurements, force data were first downsampled to 15 Hz using a moving mean to match distance measurements. Then, rupture forces were found by using the first derivative of the force data. For lifetimes, the end of the force ramp and the terminal rupture were both reported using the first derivative of the force data. For lifetimes, data were downsampled further to 5 Hz, which was necessary to automate detection of the last point of the force ramp. Then, lifetimes were calculated by taking the difference between the two timepoints and the force was averaged over the duration of each lifetime.

After finding rupture forces, plotted data were fit to the Evans-Ritchie (33) and Dudko-Hummer-Szabo (34) models for molecular adhesions to extract the intrinsic time constant and distance to transition state. Where τ is the intrinsic lifetime, r is the loading rate, and x^‡^ is the distance to transition state. From these fits, we also calculated the most probable rupture forces (F^*^). We fit the distance to transition state globally for both models as it varied little with the loading rate.

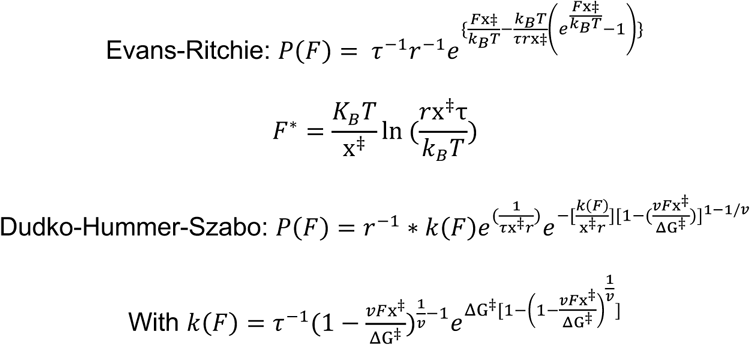

### Biochemical degradation assays

FITC-Casein (Pierce™ Fluorescent Protease Assay Kit, Thermo Scientific) was prepared as directed to 5 mg/mL in ddH_2_ O and stored at −20°C. For experiments, reactions were made in PD buffer without FITC-Casein and incubated at 30°C for >10 minutes. After incubation, FITC-Casein was added to final concentration of 0.1 mg/mL and 50 μL reactions were pipetted into a 384 well plate (Greiner Bio-One). The fluorescence was tracked using a Biotek Syngergy HTX multi-mode plate reader (Agilent Technologies) every 30 seconds with excitation/emission wavelengths of 502/528 nm. Similarly, for CM-titin degradation, reactions were made in PD buffer without substrate and incubated at 30°C for >10 minutes, after which CM-titin was added to a final concentration of 2 μM and time started. Each time point taken was quenched with a final concentration of 2x SDS-PAGE assay buffer and flash frozen in liquid nitrogen. Samples were boiled at 95°C for 5 minutes and 12% SDS-PAGE followed by staining with Coomassie Brilliant Blue.

## RESULTS

### Single-molecule mechanics of the ClpP peptidase

To probe the single-molecule mechanics of ClpP engaging an unfolded protein substrate, we used a dual-laser optical trap in passive mode (Figs. 1 B and C) without force feedback to maintain constant force (35). We immobilized biotinylated ClpP to one streptavidin-coated bead and a model multidomain substrate to separate streptavidin-coated beads. The substrate comprised a HaloTag domain at its N-terminus, which was conjugated to a biotinylated 3500bp DNA linker, tandem repeats of a variant of the I27 domain of human titin (titin^I27^) that were chemically denatured by <u>c</u>arboxy<u>m</u>ethylating buried cysteine residues (CM-titin), and a C-terminal ssrA degron tag. For optical trapping experiments, we used a microfluidic device to introduce ClpP and substrate beads into a flow cell for staged assembly in laminar flow of the ClpP-substrate complex in the absence and presence of saturating concentrations of ADEP1 (Fig. 1 C). Since tethers rarely formed in the absence of ADEP1, ADEP1 was present in all optical trapping experiments.

First, we measured the lifetimes of the interaction between ClpP and substrate at constant force at various applied loads (Fig. 2 A). The interactions between ClpP and substrate were extremely stable and did not show any translocation during the experimental timecourse, up to 200-300 seconds. The distribution of lifetimes as a function of applied load followed a slip bond behavior and fit to the Bell model of force-dependent bond rupture between two molecules separated by a potential barrier (36) (Fig. 2 B). Specifically, we were interested in measuring the unloaded lifetime (τ_0_ =1/k) and the distance to the transition state (x^‡^). From this fit, we obtained an unloaded lifetime (τ_0_) of 158 ± 39 s and a distance to the transition state (x^‡^) of 0.3 ± 0.1 nm (Mean ± SEM, N=43). We note that this unloaded lifetime is much longer than the unfolding time constants of several substrate domains from single-molecule studies of ClpAP and ClpXP, which vary between 0.3-55 s for several variants of titin^I27^, 0.03-3.4 s for filamin domains, and 9.1-19 s for GFP (5–8, 12, 17, 37).

**FIGURE 2.**
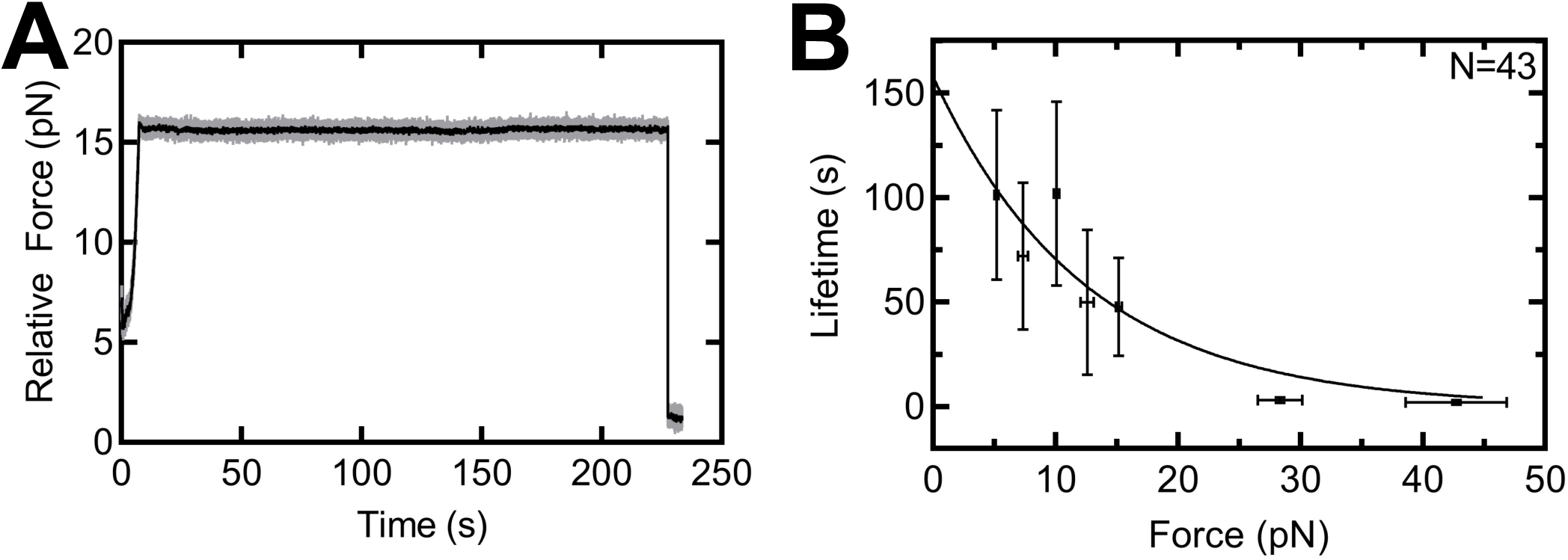
ADEP-ClpP forms long lived interactions with protein substrate under load. *(A)* Example time course of ClpP interaction with CM-titin as a function of applied load. Data were downsampled to 700 Hz (gray) and 50 Hz (black). A constant speed force ramp was applied until the target force reached, after which the trap position remained constant until tether rupture back to 0 pN. *(B)* Tether lifetimes as a function of applied load showing the mean ± SEM in both x and y (N=8, 4, 8, 2, 9, 3 for each force from low to high). The solid line is the fit to the Bell-Evans model for a slip bond (see Methods) yielding τ = 158 ± 39 s and x^‡^ = 0.3 ± 0.11 nm (fit ± SEM).

Because of the observed long timescales of the experiments above and lack of translocation, we also measured rupture force as a function of loading rate to probe the mechanical strength of the interaction between ClpP and substrate. A linear force ramp was applied to the ClpP-substrate tether until a terminal rupture occurred (Fig. 3 A). We observed a bimodal behavior in ClpP-substrate interactions as demonstrated from the distributions of rupture forces under different loading rates (Fig. 3 B). To exclude the possibility of artifacts arising from tethering our enzyme-substrate complex via a DNA linker or from the experimental dual bead geometry, we examined the force-induced rupture of an oligo annealed to DNA to ensure the validity of our dual-bead assay. Our results using the same 3500bp DNA linker with a 20bp overhang without ligation agreed with previously published literature on DNA shearing (38), yielded different rupture forces than any of the rupture peaks observed for ClpP-substrate tethers, and did not display bimodal behavior, giving us confidence in our experimental design (Fig. S1).

**FIGURE 3.**
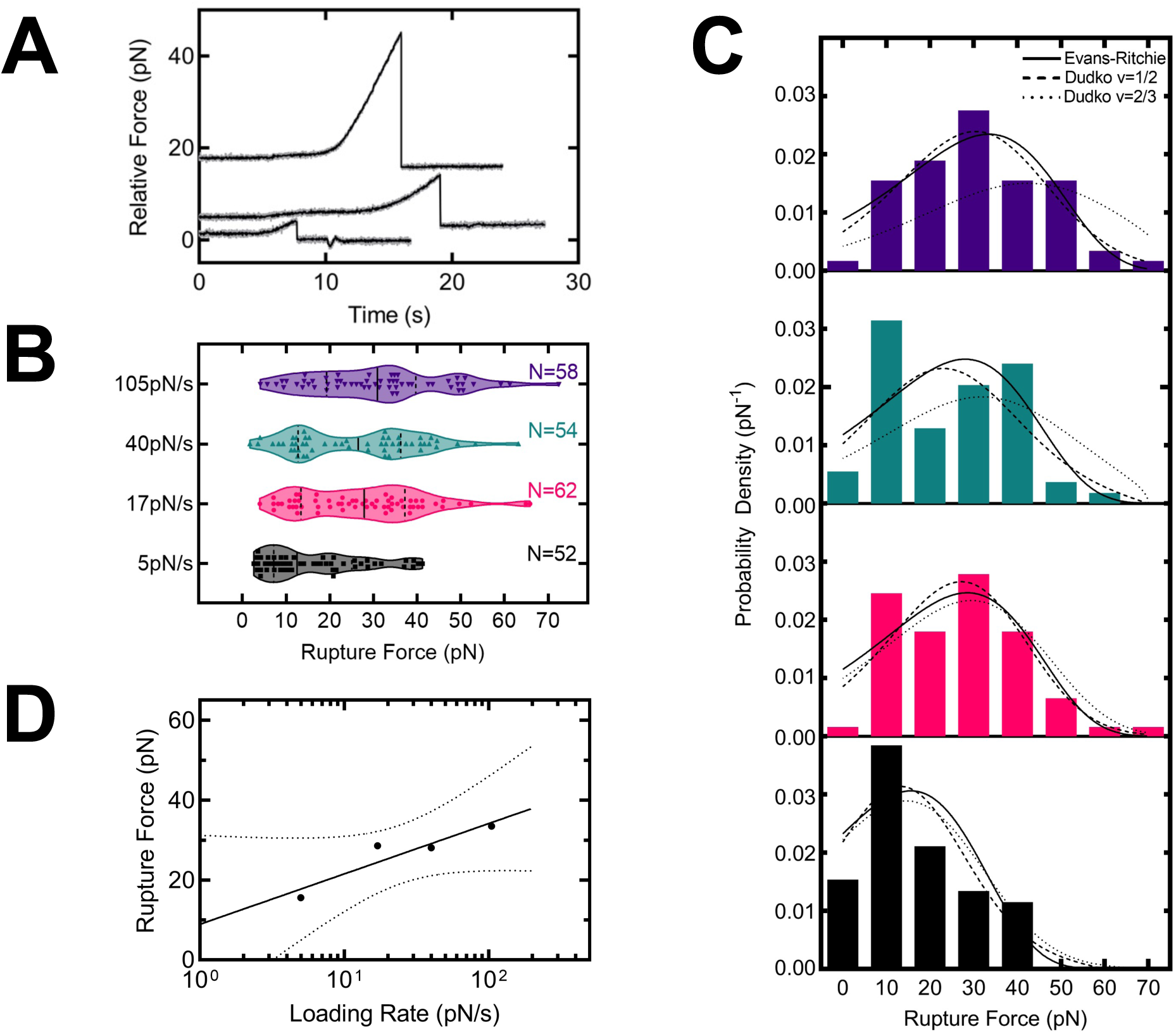
ADEP-ClpP grips substrate against external load. *(A)* Example rupture force traces of ADEP-ClpP engaging CM-titin. Data were downsampled to 700 Hz (gray) and 50 Hz (black). Example traces are offset on the y-axis for clarity. A constant speed force ramp is applied until the interaction ruptures to 0 pN. *(B)* Violin plots of ClpP-substrate rupture forces in the presence of ADEP are shown at indicated loading rates. Data points represent unique tethers with a terminal rupture to 0 pN. Vertical lines mark the median and quartiles of each distribution and N of each loading rate is shown. *(C)* Histograms of the rupture forces shown in *(B)*. Fits to the Evans-Ritchie model (33) for each loading rate are shown as solid black lines with v=1/2 (dashed lines) and v=2/3 (dotted lines) for Dudko-hummer-Szabo (34) fits. *(D)* Most probable rupture force is plotted as a function of loading rate for wild type ClpP-substrate interactions in the presence of ADEP. The most probable rupture forces shown are derived from fits of data in *(C)* to the Evans-Ritchie model. The data were fit to a semi log line with parameters b = 7.6 ± 7.3 pN and m = 13.4 ± 4.9 (Fit +/- SEM).

For ClpP-substrate interactions, the rupture force distributions at different loading rates were fit to several models of bond rupture based on Kramer’s Theory, the Evans-Ritchie (33) and Dudko-Hummer-Szabo (34) models (Fig 3 C). From these fits, we obtained the thermal off rate (*k*), distance to the transition state (x^‡^), and the free energy of activation (ΔG^‡^) for the ClpP-substrate complex in the presence of ADEP1 (Table 1). For the Evans Ritchie model, the fits for the unloaded lifetimes (τ_0_ =1/k) varied between 1-10 s depending on the loading rate, though this variability may represent the crossing of an energy barrier as seen for other molecular interactions (39, 40). The Dudko-Hummer-Szabo model takes the shape of the transition state surface into account, which can be either a cusp (v=1/2) or linear-cubic (v=2/3). The (τ_0_) of these fits were similar to the Evans Ritchie parameters while the (x^‡^) was similar for the linear-cubic and slightly larger for the cusp profile, 0.3 ± 0.1 nm and 0.4 ± 0.1 nm, respectively (mean ± SEM). The free energy of activation varied with the loading rate yielding values between 3-5 k_B_ T (Table 1). Finally, we fit the most probable rupture forces calculated from the Evans Ritchie model as a function of loading rate to a semi-log line with intercept (b) = 8.9 ± 5.2 pN and slope (m) = 12.6 ± 3.5 (Fig 3 D). This slope represents the force sensitivity of the interaction as it is the force necessary to increase the dissociation rate *k*_off_ by *e*-fold.

**TABLE 1.** Fits for rupture force distributions shown in Figures 3 and 4. Table of fit parameters to the Evans-Ritchie and Dudko-Hummer-Szabo models (see Methods). Values shown are the best fit ± SEM.

| Evans-Ritchie Model Parameters |  | wild type ClpP |  | ClpPS97A |  |  |
| --- | --- | --- | --- | --- | --- | --- |
| Loading Rates (pN/s) | $\tau$ (s) | $x^\ddagger$ (nm) | | $\tau$ (s) | $x^\ddagger$ (nm) | |
| 5 | $8.59 \pm 1.44$ | 0.22± 0.03 | | $9.68 \pm 2.11$ | $0.25 \pm 0.04$ | |
| 17 | $5.10 \pm 1.16$ | | | $6.60 \pm 1.99$ | | |
| 40 | $2.11 \pm 0.47$ | | | $2.90 \pm 0.88$ | | |
| 105 | $1.08 \pm 0.27$ | | | $1.22 \pm 0.38$ | | |
| Dudko-Hummer-Szabo Parameters $\nu = 1/2$ | | wild type ClpP | | ClpPS97A | | |
| Loading Rates (pN/s) | $\tau$ (s) | $x^\ddagger$ (nm) | $\Delta G$ (kT) | $\tau$ (s) | $x^\ddagger$ (nm) | $\Delta G$ (kT) |
| 5 | $9.16 \pm 1.90$ | $0.37 \pm 0.07$ | $2.74 \pm 0.57$ | $10.49 \pm 3.04$ | $0.41 \pm 0.10$ | $3.19 \pm 1.13$ |
| 17 | $6.88 \pm 2.06$ | | $4.73 \pm 2.13$ | $9.67 \pm 4.27$ | | $5.47 \pm 3.06$ |
| 40 | $2.43 \pm 0.77$ | | $3.19 \pm 0.71$ | $3.41 \pm 1.66$ | | $3.19 \pm 0.95$ |
| 105 | $1.42 \pm 0.48$ | | $4.43 \pm 1.59$ | $1.56 \pm 0.72$ | | $4.29 \pm 1.67$ |
| Dudko-Hummer-Szabo Parameters $\nu= 2/3$ | | wild type ClpP | | ClpPS97A | | |
| Loading Rates (pN/s) | $\tau$ (s) | $x^\ddagger$ (nm) | $\Delta G$ (kT) | $\tau$ (s) | $x^\ddagger$ (nm) | $\Delta G$ (kT) |
| 5 | $8.97 \pm 2.26$ | $0.25 \pm 0.07$ | $2.93 \pm 3.39$ | $9.34 \pm 2.91$ | $0.26 \pm 0.07$ | $4.06 \pm 8.33$ |
| 17 | $5.89 \pm 2.17$ | | $4.95 \pm 7.33$ | $7.48 \pm 4.33$ | | Unstable |
| 40 | $3.22 \pm 1.56$ | | $2.93 \pm 0.82$ | $3.92 \pm 2.34$ | | $2.96 \pm 0.74$ |
| 105 | $2.24 \pm 1.43$ | | $3.18 \pm 0.86$ | $2.78 \pm 2.14$ | | $3.10 \pm 0.81$ |

### Contribution of the ClpP active site to substrate grip

Having observed the ability of ClpP to grip protein substrate under load, we asked if we could determine what domain of ClpP contributes to the peptidase’s mechanical behavior. We hypothesized that the ClpP active site might affect substrate grip through formation of a covalent intermediate, as observed for all serine proteases, by coordinating substrate binding, or through a combination of both mechanisms. Therefore, we first mutated the active site serine to alanine (ClpPS97A) to assess how the active site affects substrate grip. ClpPS97A inactivation was verified by monitoring the degradation of a fluorescently-labeled unfolded substrate (FITC-Casein) and of our model CM-titin substrate (Fig. S2). In rupture force experiments, the shape of ClpPS97A distributions remained similar to wild type ClpP (Fig. 4 A). We fit the distribution of rupture forces to the Evans-Ritchie and Dudko-Hummer-Szabo models, which yielded similar off rates, distance to the transition state, and free energy of activation as wild type ClpP (Fig. 4 B and Table 1). Finally, we fit the most probable rupture forces as a function of loading rate for ClpPS97A and found that the slope and intercept also remained similar to wild type ClpP. These data suggest that the ClpP active site does not contribute to substrate grip. In addition to the active site mutation, we used diisopropylfluorophosphate (DFP) to verify if the ClpP active site contributes to grip, as DFP chemically inactivates ClpP. We found that the shape of the distribution still remained similar to both wild type ClpP and ClpPS97A at the tested loading rate (Fig. S2). This further supports the conclusion that the ClpP active site is not coupled to mechanical gripping of protein substrate. Using the parameters from the Dudko-Hummer-Szabo model, we recreated the transition state energy landscapes of the observed ClpP-substrate interactions with energy wells represented as harmonic potentials (Fig. 5). We compared these to other measured protein-protein interactions (41) and found that the free energy of activation and distance to the transition state are comparable to the interaction between fluorescein and an anti-fluorescein antibody. Interestingly, the transition state distance is also similar (∼0.7-1 nm) to those observed for force-dependent ClpXP translocation along substrate polypeptide (5, 11).

**FIGURE 4.**
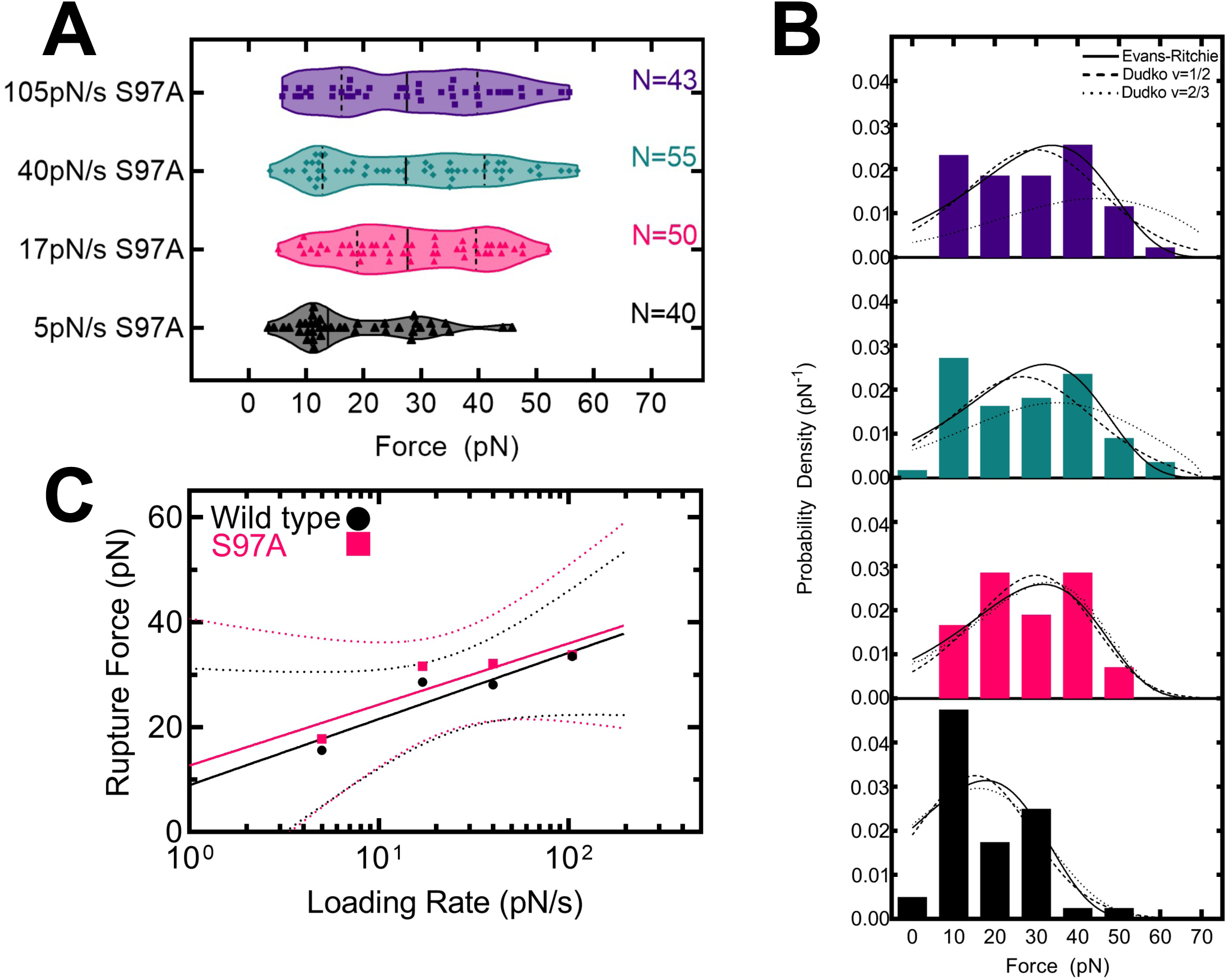
ClpP active site inactivation does not affect substrate grip. *(A)* Violin plots of ClpPS97A-substrate rupture forces in the presence of ADEP are shown at indicated loading rates. Data points represent unique tethers with a terminal rupture to 0 pN. Vertical lines mark the median and quartiles of each distribution. *(B)* Histograms of the rupture forces shown in *(A)*. Fits to the Evans-Ritchie model for each loading rate are shown as solid black lines while Dudko-Hummer-Szabo fits with v=1/2 (dashed lines) and v=2/3 (dotted lines). *(C)* Most probable rupture force is plotted as a function of loading rate for wild type ClpP- and ClpPS97A-substrate interactions in the presence of ADEP. The most probable rupture forces are derived from fits of data in *(B)* to the Evans-Ritchie model. For ClpPS97A, the data was fit to a semi log line with parameters b= 13.9 ± 6.4 pN and m = 10.5 +/- 4.3 (Fit +/- SEM).

**FIGURE 5.**
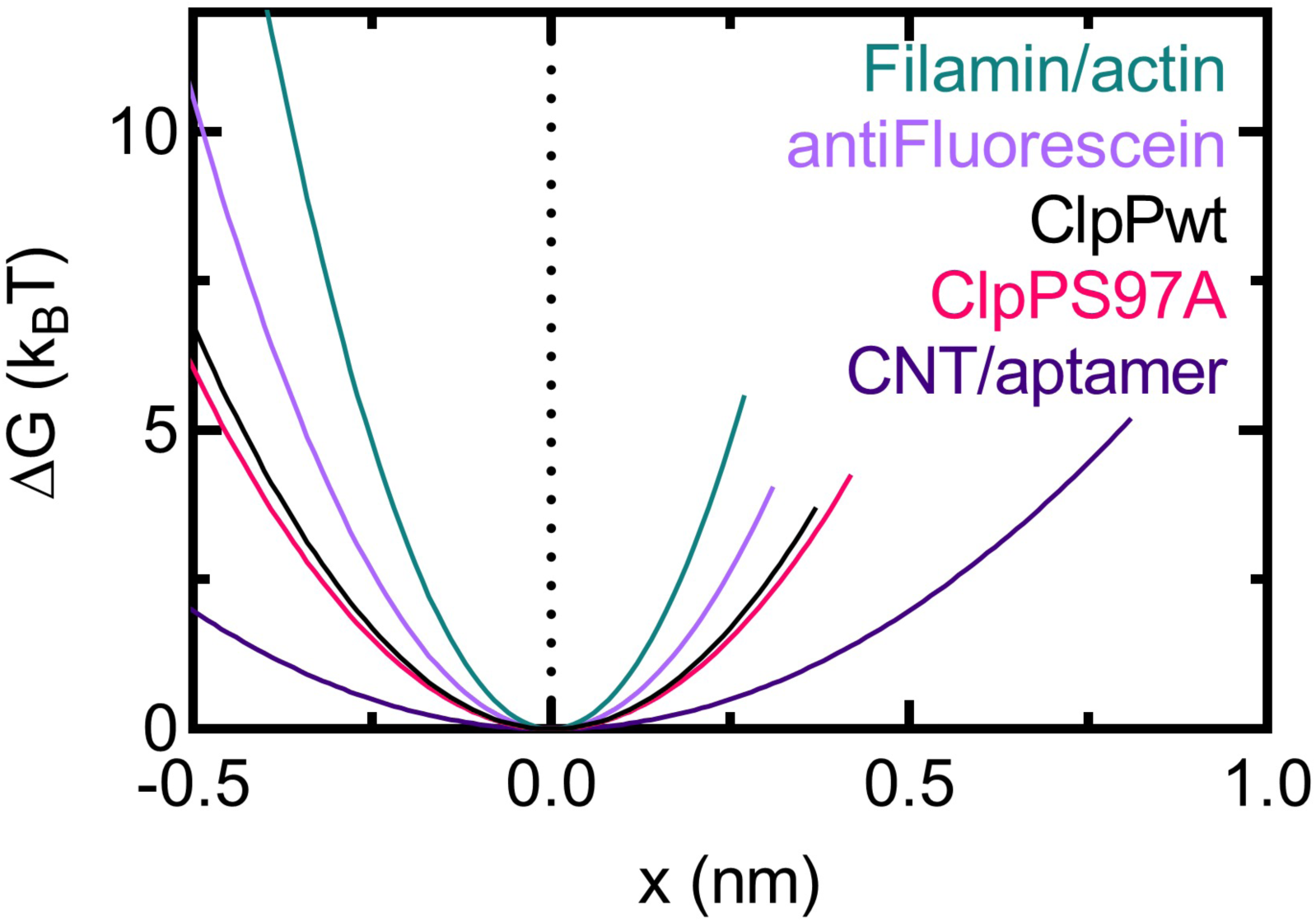
Free energy diagram of the ClpP-substrate interactions. Free energy diagrams were constructed according to the Dudko-Hummer-Szabo model (34) assuming a cusp shape (v=1/2). Each curve represents a different interaction graphed in x until the distance to the transition state, at which point the curve stops. Both wild type ClpP and ClpPS97A show a similar energy barrier to antifluorescein-fluorescein. Values for several other interactions were taken from Refs. (41, 63) for comparison.

## DISCUSSION

Protein degradation by AAA+ proteases is an essential cellular process that requires the coordination of ATP-dependent motor proteins with their peptidase components. Based on single-molecule studies (1, 2, 5–8, 10, 11), the AAA+ proteases ClpXP and ClpAP produce pN forces to unfold and translocate protein substrates. The motors grip substrate using distinct pore loops (10, 11, 17) and drive polypeptide translocation through conformational changes between motor subunits (7, 8, 42). However, whether the peptidase, ClpP, aids mechanically to this reaction remained a question of interest. Here, using single-molecule optical trapping, we show the first evidence that ClpP grips an unfolded protein substrate against significant external load, in the absence of a AAA+ motor. The force-dependent lifetimes follow characteristic slip bond behavior (Fig. 2), with an average unloaded lifetime of 157 seconds. Notably, this is similar to ClpAP and ClpXP-mediated unfolding lifetimes of difficult-to-unfold substrates like the wild type titin^I27^ domain (5–8, 10, 12, 17, 37) and helps explain the trapping of substrates by proteolytically inactive ClpP variants in proteomic studies (43–45). We hypothesize that the observed force-dependent interaction between ClpP and substrate contributes to overall substrate grip during AAA+ protease degradation and likely aids in preventing slipping of protein substrates. Our hypothesis is consistent with experimental data demonstrating that addition of an unfolded region prior to the folded domain of GFP-ssrA increases the unfolding and degradation speed by ClpXP (46), that slipping events are readily observed during unfolding and translocation of substrate by ClpX in the absence of ClpP (5, 6, 10), and that a longer unstructured substrate tail capable of reaching into ClpP compensates for reduced substrate grip by pore 1-loop variants of ClpXP (10).

Furthermore, we characterized ClpP-substrate rupture forces and fit them to models of force-dependent protein-ligand interaction based on Kramers theory (33, 34). The fits yield small distances to the transition state, (x^‡^=0.2-0.3nm) while unloaded bond lifetimes (τ=1/k_0_) vary between 1-10s depending on the loading rate (Fig. 3, Table 1). Based on these parameters, we obtain a free energy of ClpP-substrate interaction, Α’G≈4 k_B_ T, which is similar to estimates of work (∼5 k_B_ T) produced by ClpXP and ClpAP during a power stroke (5, 12). Therefore, the energetics contributing to substrate grip by ClpP could provide a partial failsafe for the AAA+ protease to remain bound to a difficult to unfold substrate or under conditions of limiting ATP concentration, such as during stationary phase in bacteria (47). During these periods of slow growth due to unfavorable conditions such as nutrient limitation, proteins could evade degradation by refolding and releasing before the motor has a chance to unfold and translocate (48).

Interestingly, we find that rupture forces distribute in a bimodal fashion. Several hypotheses account for bimodality. First, at least two populations arise from interactions of the substrate with distinct ClpP conformers, such as those observed in structural studies (49–53). Second, ClpP possesses two substrate binding regions or sites, each responding to external force differently. We conjectured that one site would be the active site since the protease reaction proceeds through a covalent intermediate (Fig 4). Here we show that mutating the active site serine to alanine did not significantly affect substrate grip by ClpP. Third, the N-terminal loops of ClpP present another candidate site mediating substrate grip, as they already play a role in substrate gating (20) and modulate the activity of the active site serine (21). Likewise, different populations could arise due to a combination of multiple ClpP subunits engaging the substrate. Finally, the bimodality could be caused by the substrate’s conformation as it enters the chamber of ClpP. For example, unfolded CM-titin possibly enters as or forms partially folded intermediates within ClpP that require greater force to rupture. Such structures are capable of being degraded by ClpXP as previous studies show that two polypeptide chains linked by a disulfide bond (4) and knotted protein substrates are degraded by ClpXP (54–56). Ultimately, ClpP grips protein substrate in the absence of a AAA+ motor protein, although the molecular details defining ClpP grip require further study.

Substrate grip exhibited by ClpP might have important implications for ClpAP and ClpXP. For example, ClpA’s unfolding speed increases when in complex with ClpP in a manner not fully accounted for by ClpA’s ATPase activation when ClpP is present, i.e. 7-fold unfolding speed increase yet 2-fold ATPase increase with ClpP (14). Additionally, a ClpX variant with unfolding defects is rescued when in complex with wild type ClpP, ClpPS97A, and DFP-labeled ClpP (57). Furthermore, ClpX slips back farther and more frequently in the absence of ClpP at the single-molecule level (5, 6, 10, 12). While the exact mechanisms are unclear, we hypothesize that substrate grip provided by ClpP helps ClpA and ClpX unfold substrate and suppress back slips by ClpX. Our data suggest that ClpP plays a more active role in degradation by aiding in substrate grip that may result in increased degradation efficiency or unfolding speed by preventing reversible folding and premature release. Likewise, the abililty of ClpP to grip protein substrates would be predicted to enhance processivity of these proteolytic enzymes by decreasing the probability of substrate release.

Processive degradation by AAA+ proteases prevents the release of partially degraded products that would be detrimental to cellular function, since these products could bind and inhibit protein partners or lead to aggregation and cell death. However, many AAA+ enzymes are weakly or not processive and do not need to be to fulfill their cellular functions. For example, spastin and katanin only partially unfold tubulin dimers to sever microtubules (58), NSF disassembles SNARE complexes without entirely unfolding and translocating individual SNARE proteins (59), and mitochondrial ClpX remodels the heme biosynthetic enzyme ALAS through partial unfolding (60). Despite similarities in structure among the various AAA+ unfoldases characterized to date (61), AAA+ proteases must possess some unique property in order to ensure processive translocation and degradation of substrates. While many studies of ClpXP and ClpAP focus on how the motor contributes to processivity, we propose that the combination of ATPase and peptidase makes a complete processive machine. The ability of the peptidase to perhaps act as a processivity factor is likely a general feature of AAA+ proteases as results from Classen et al. show that the proteasomal 20S core particle also maintains protein substrate grip under load (62). However, the 20S active site threonine contributes to the observed mechanical behavior of the enzyme, which we do not observe here for the homologous ClpP active site serine. Therefore, there are likely key differences to how the peptidase components of AAA+ proteases contribute to overall mechanical degradation.

## AUTHOR CONTRIBUTIONS

S.W. and A.O. designed research; S.W. performed research; S.W. contributed analytical tools; S.W and A.O. analyzed data; S.W. and A.O wrote the manuscript.

## ACKNOWLEDGEMENTS

We thank Jennifer Norton and members of the Olivares and Lang laboratories for insightful discussions related to the hypothesis of the present work. We also thank Marija Zanic, Matt Lang and Gregor Neuert for critical feedback on the manuscript and Manny Ascano for use of his plate reader. This work was supported by start-up funds provided by the Department of Biochemistry at Vanderbilt University and in part by training grant T32GM008320 (NIGMS, NIH).

---

**FIGURE S1.**
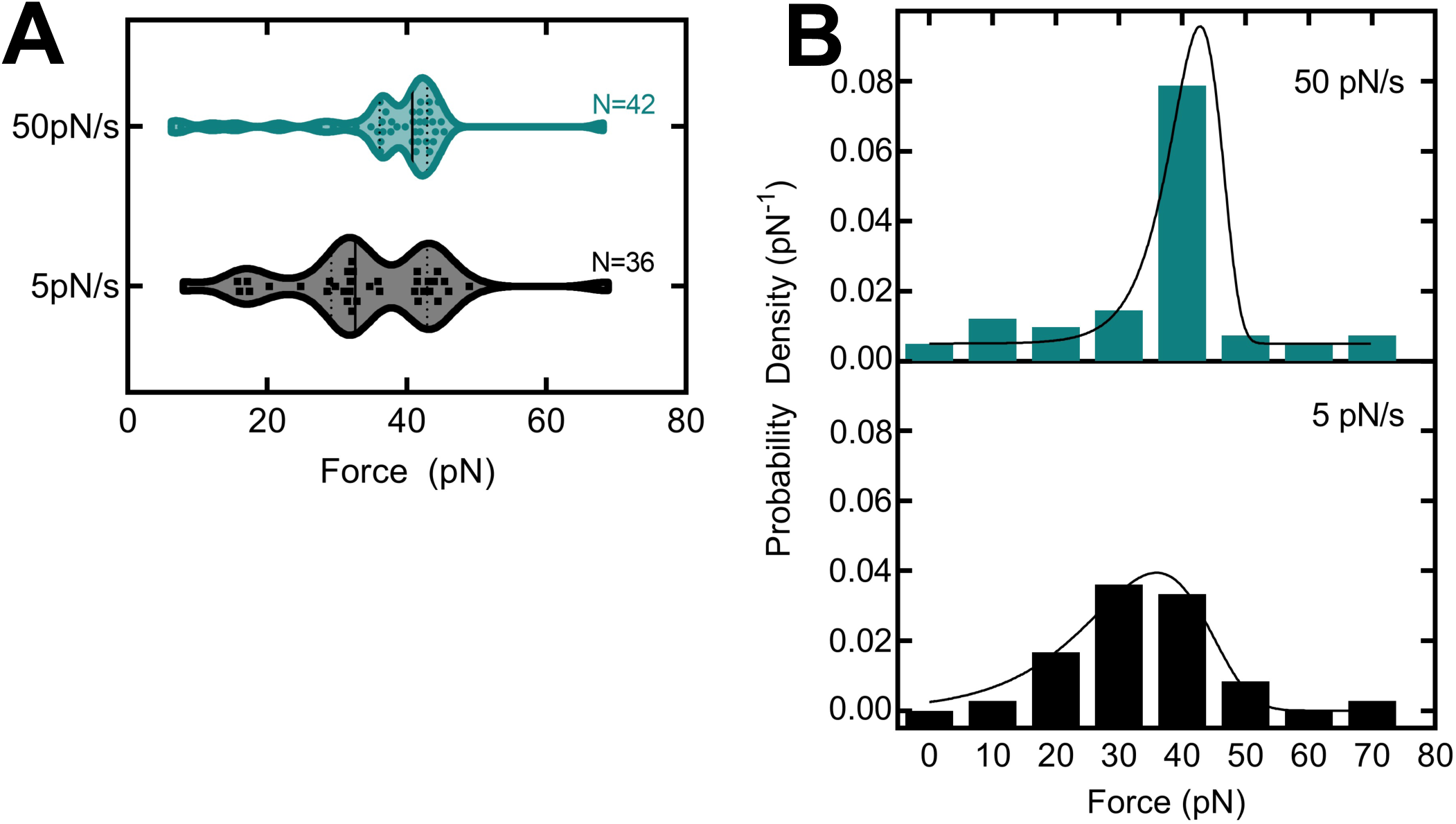
Controlling for DNA overhang effects on observed rupture forces. *(A)* Violin plots of observed rupture forces for control linker DNA with overhang at two different loading rates (see Methods). Data points represent unique tethers with a terminal rupture to 0 pN. Vertical lines mark the median and quartiles of each distribution. *(B)* Histograms of the rupture forces shown in *(A)*. Fits to the Evans-Ritchie model for each loading rate are shown as solid black lines.

**FIGURE S2.**
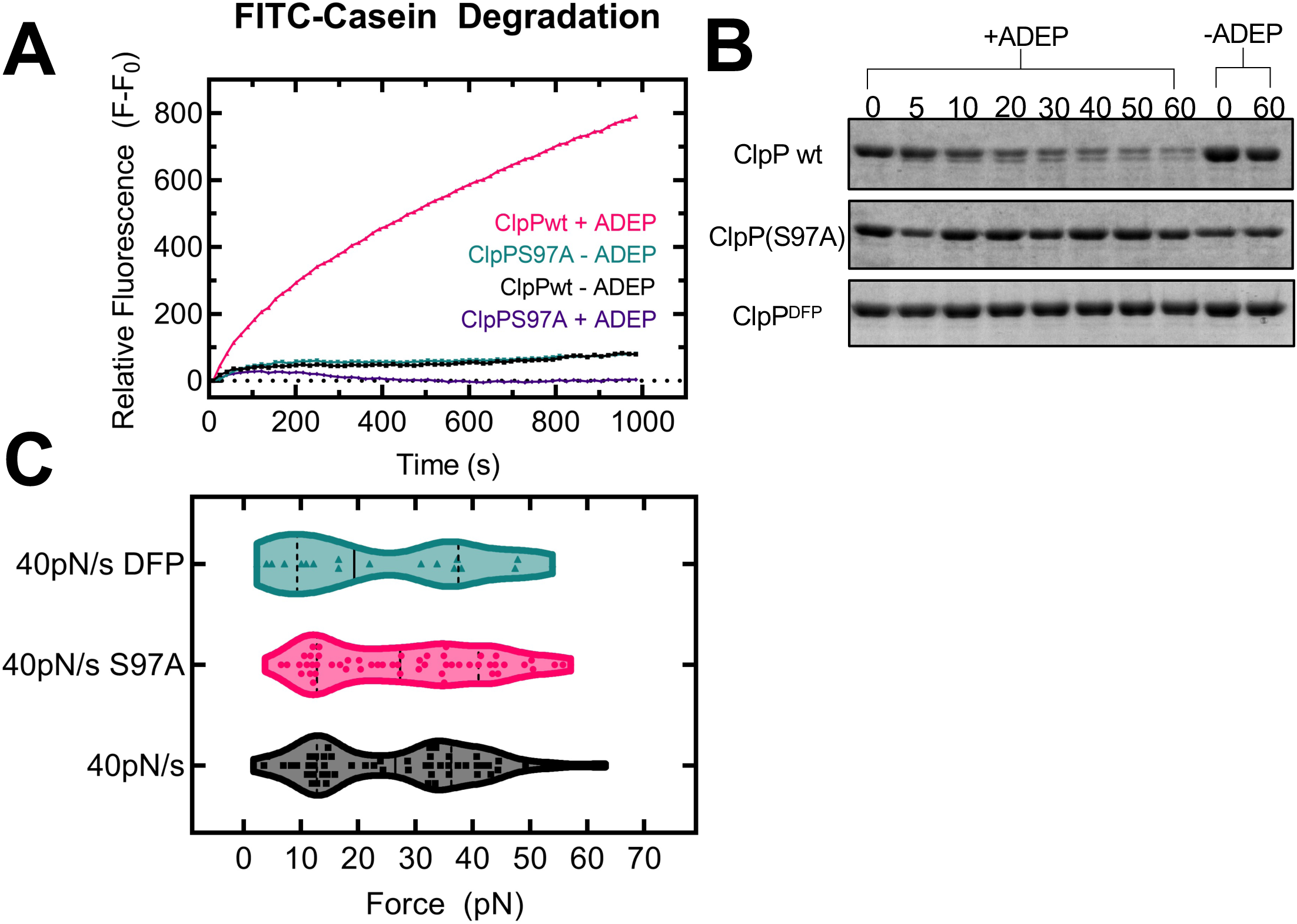
Mutation of Ser97 to Ala and DFP inactivate ClpP for degradation. *(A)* FITC-Casein degradation assay measuring proteolysis by ClpP. As FITC-Casein is degraded, FITC fluorescence increases over time. Only wild type ClpP is active for degradation in the presence of ADEP over this timecourse (N=1). *(B)* Degradation of the multidomain CM-titin substrate monitored by SDS-PAGE. ClpPS97A and DFP-ClpP do not easily degrade this substrate in comparison to wild type ClpP. C) Violin plots of rupture forces comparing wild type ClpP, ClpPS97A, and DFP-ClpP at 40 pN/s loading rate. The median and quartiles are represented by the solid and dashed lines, respectively.

## REFERENCES

1. Olivares, A.O., T.A. Baker, and R.T. Sauer. 2018. Mechanical Protein Unfolding and Degradation. Annu. Rev. Physiol. 80:413–429.

2. Olivares, A.O., T.A. Baker, and R.T. Sauer. 2015. Mechanistic insights into bacterial AAA+ proteases and protein-remodelling machines. Nat. Rev. Microbiol. 14:33–44.

3. Maurizi, M.R., W.P. Clark, S.H. Kim, and S. Gottesman. 1990. Clp P represents a unique family of serine proteases. J. Biol. Chem. 265:12546–12552.

4. Burton, R.E., S.M. Siddiqui, Y.I. Kim, T.A. Baker, and R.T. Sauer. 2001. Effects of protein stability and structure on substrate processing by the ClpXP unfolding and degradation machine. EMBO J. 20:3092–3100.

5. Aubin-Tam, M.-E., A.O. Olivares, R.T. Sauer, T.A. Baker, and M.J. Lang. 2011. Single-Molecule Protein Unfolding and Translocation by an ATP-Fueled Proteolytic Machine. Cell. 145:257–267.

6. Maillard, R.A., G. Chistol, M. Sen, M. Righini, J. Tan, C.M. Kaiser, C. Hodges, A. Martin, and C. Bustamante. 2011. ClpX(P) generates mechanical force to unfold and translocate its protein substrates. Cell. 145:459–469.

7. Sen, M., R.A. Maillard, K. Nyquist, P. Rodriguez-Aliaga, S. Pressé, A. Martin, and C. Bustamante. The ClpXP protease unfolds substrates using a constant rate of pulling but different gears. Cell. 155:636–646.

8. Cordova, J.C., A.O. Olivares, Y. Shin, B.M. Stinson, S. Calmat, K.R. Schmitz, M.E. Aubin-Tam, T.A. Baker, M.J. Lang, and R.T. Sauer. 2014. Stochastic but highly coordinated protein unfolding and translocation by the ClpXP proteolytic machine. Cell. 158:647–658.

9. Olivares, A.O., A.R. Nager, O. Iosefson, R.T. Sauer, and T.A. Baker. 2014. Mechanochemical basis of protein degradation by a double-ring AAA+ machine. Nat. Struct. Mol. Biol. 21:871–875.

10. Iosefson, O., A.O. Olivares, T.A. Baker, and R.T. Sauer. 2015. Dissection of axial-pore loop function during unfolding and translocation by a AAA+ proteolytic machine. Cell Rep. 12:1032– 1041.

11. Rodriguez-Aliaga, P., L. Ramirez, F. Kim, C. Bustamante, and A. Martin. 2016. Substrate-translocating loops regulate mechanochemical coupling and power production in AAA+ protease ClpXP. Nat. Struct. & Mol. Biol. 23:974–982.

12. Olivares, A.O., H.C. Kotamarthi, B.J. Stein, R.T. Sauer, and T.A. Baker. 2017. Effect of directional pulling on mechanical protein degradation by ATP-dependent proteolytic machines. Proc. Natl. Acad. Sci. 114:E6306 LP–E6313.

13. Kotamarthi, H.C., R.T. Sauer, and T.A. Baker. 2020. The Non-dominant AAA+ Ring in the ClpAP Protease Functions as an Anti-stalling Motor to Accelerate Protein Unfolding and Translocation. Cell Rep. 30:2644-2654.e3.

14. Baytshtok, V., T.A. Baker, and R.T. Sauer. 2015. Mechanism of protein remodeling by ClpA chaperone. Proc. Natl. Acad. Sci. 112:5377–5382.

15. Rajendar, B., and A.L. Lucius. 2010. Molecular Mechanism of Polypeptide Translocation Catalyzed by the Escherichia coli ClpA Protein Translocase. J. Mol. Biol. 399:665–679.

16. Miller, J.M., J. Lin, T. Li, and A.L. Lucius. 2013. E. coli ClpA catalyzed polypeptide translocation is allosterically controlled by the protease ClpP. J. Mol. Biol. 425:2795–2812.

17. Iosefson, O., A.R. Nager, T.A. Baker, and R.T. Sauer. 2015. Coordinated gripping of substrate by subunits of a AAA+ proteolytic machine. Nat. Chem. Biol. 11:201–206.

18. Hinnerwisch, J., B.G. Reid, W.A. Fenton, and A.L. Horwich. 2005. Roles of the N-domains of the ClpA unfoldase in binding substrate proteins and in stable complex formation with the ClpP protease. J. Biol. Chem. 280:40838–40844.

19. Kim, Y.I., I. Levchenko, K. Fraczkowska, R. V. Woodruff, R.T. Sauer, and T.A. Baker. 2001. Molecular determinants of complex formation between Clp/Hsp 100 ATPases and the ClpP peptidase. Nat. Struct. Biol. 8:230–233.

20. Gribun, A., M.S. Kimber, R. Ching, R. Sprangers, K.M. Fiebig, and W.A. Houry. 2005. The ClpP double ring tetradecameric protease exhibits plastic ring-ring interactions, and the N termini of its subunits form flexible loops that are essential for ClpXP and ClpAP complex formation. J. Biol. Chem. 280:16185–16196.

21. Jennings, L.D., J. Bohon, M.R. Chance, and S. Licht. 2008. The ClpP N-terminus coordinates substrate access with protease active site reactivity. Biochemistry. 47:11031–11040.

22. Brötz-Oesterhelt, H., D. Beyer, H.P. Kroll, R. Endermann, C. Ladel, W. Schroeder, B. Hinzen, S. Raddatz, H. Paulsen, K. Henninger, J.E. Bandow, H.G. Sahl, and H. Labischinski. 2005. Dysregulation of bacterial proteolytic machinery by a new class of antibiotics. Nat. Med. 11:1082– 1087.

23. Malik, I.T., and H. Brötz-Oesterhelt. 2017. Conformational control of the bacterial Clp protease by natural product antibiotics. Nat. Prod. Rep. 34:815–831.

24. Amor, A.J., K.R. Schmitz, J.K. Sello, T.A. Baker, and R.T. Sauer. 2016. Highly Dynamic Interactions Maintain Kinetic Stability of the ClpXP Protease during the ATP-Fueled Mechanical Cycle. ACS Chem. Biol. 11:1552–1560.

25. Gersch, M., K. Famulla, M. Dahmen, C. Göbl, I. Malik, K. Richter, V.S. Korotkov, P. Sass, H. Rübsamen-Schaeff, T. Madl, H. Brötz-Oesterhelt, and S.A. Sieber. 2015. AAA+ chaperones and acyldepsipeptides activate the ClpP protease via conformational control. Nat. Commun. 6.

26. Conlon, B.P., E.S. Nakayasu, L.E. Fleck, M.D. Lafleur, V.M. Isabella, K. Coleman, S.N. Leonard, R.D. Smith, J.N. Adkins, and K. Lewis. 2013. Activated ClpP kills persisters and eradicates a chronic biofilm infection. Nature. 503:365–370.

27. Graves, P.R., L.J. Aponte-Collazo, E.M.J. Fennell, A.C. Graves, A.E. Hale, N. Dicheva, L.E. Herring, T.S.K. Gilbert, M.P. East, I.M. McDonald, M.R. Lockett, H. Ashamalla, N.J. Moorman, D.S. Karanewsky, E.J. Iwanowicz, E. Holmuhamedov, and L.M. Graves. 2019. Mitochondrial Protease ClpP is a Target for the Anticancer Compounds ONC201 and Related Analogues. ACS Chem. Biol. 14:1020–1029.

28. Moreno-cinos, C., K. Goossens, I.G. Salado, and P. Van Der Veken. 2019. ClpP Protease, a Promising Antimicrobial Target. Int. J. Mol. Sci. 20.

29. Sass, P., M. Josten, K. Famulla, G. Schiffer, H.-G. Sahl, L. Hamoen, and H. Brötz-Oesterhelt. Antibiotic acyldepsipeptides activate ClpP peptidase to degrade the cell division protein FtsZ. Proc. Natl. Acad. Sci. 108:17474–17479.

30. Silber, N., S. Pan, S. Schä;kermann, C. Mayer, H. Brötz-Oesterhelt, and P. Sass. 2020. Cell Division Protein FtsZ Is Unfolded for N-Terminal Degradation by Antibiotic-Activated ClpP. MBio. 11:e01006–20.

31. Cordova, J.C., A.O. Olivares, and M.J. Lang. 2017. Mechanically watching the ClpXP proteolytic machinery. In: Methods in Molecular Biology. Humana Press Inc. pp. 317–341.

32. Kenniston, J.A., T.A. Baker, J.M. Fernandez, and R.T. Sauer. 2003. Linkage between ATP consumption and mechanical unfolding during the protein processing reactions of an AAA+ degradation machine. Cell. 114:511–520.

33. Evans, E., and K. Ritchie. 1997. Dynamic Strength of Molecular Adhesion Bonds. Biophys. J. 72:1541–1555.

34. Dudko, O.K., G. Hummer, and A. Szabo. 2006. Intrinsic rates and activation free energies from single-molecule pulling experiments. Phys. Rev. Lett. 96:1–4.

35. Bustamante, C.J., Y.R. Chemla, S. Liu, and M.D. Wang. 2021. Optical tweezers in single-molecule biophysics. Nat. Rev. Methods Prim. 1.

36. Bell, G.I. 1978. Models for the specific adhesion of cells to cells. Science (80-.). 200:618 LP – 627.

37. Shin, Y., J.H. Davis, R.R. Brau, A. Martin, J.A. Kenniston, T.A. Baker, R.T. Sauer, and M.J. Lang. 2009. Single-molecule denaturation and degradation of proteins by the AAA+ ClpXP protease. Proc. Natl. Acad. Sci. 106:19340–19345.

38. Lang, M.J., P.M. Fordyce, A.M. Fordyce, K.C. Neuman, and S.M. Block. 2004. Simultaneous, coincident optical trapping and single-molecule fluorescence. Nat. Methods. 1:133–139.

39. Neuert, G., C. Albrecht, E. Pamir, and H.E. Gaub. 2006. Dynamic force spectroscopy of the digoxigenin-antibody complex. FEBS Lett. 580:505–509.

40. Merkel, R., P. Nassoy, A. Leung, K. Ritchie, and E. Evans. 1999. Energy landscapes of receptor-ligand bonds explored with dynamic force spectroscopy. Nature. 397:50–53.

41. Aubin-Tam, M.E., D.C. Appleyard, E. Ferrari, V. Garbin, O.O. Fadiran, J. Kunkel, and M.J. Lang. Adhesion through single peptide aptamers. J. Phys. Chem. A. 115:3657–3664.

42. Stinson, B.M., V. Baytshtok, K.R. Schmitz, T.A. Baker, and R.T. Sauer. 2015. Subunit asymmetry and roles of conformational switching in the hexameric AAA+ ring of ClpX. Nat. Struct. & Mol. Biol. 22:411.

43. Flynn, J.M., S.B. Neher, Y.-I. Kim, R.T. Sauer, and T.A. Baker. 2003. Proteomic Discovery of Cellular Substrates of the ClpXP Protease Reveals Five Classes of ClpX-Recognition Signals. Mol. Cell. 11:671–683.

44. Neher, S.B., J. Villén, E.C. Oakes, C.E. Bakalarski, R.T. Sauer, S.P. Gygi, and T.A. Baker. 2006. Proteomic Profiling of ClpXP Substrates after DNA Damage Reveals Extensive Instability within SOS Regulon. Mol. Cell. 22:193–204.

45. Feng, J., S. Michalik, A.N. Varming, J.H. Andersen, D. Albrecht, L. Jelsbak, S. Krieger, K. Ohlsen, M. Hecker, U. Gerth, H. Ingmer, and D. Frees. 2013. Trapping and proteomic identification of cellular substrates of the ClpP protease in staphylococcus aureus. J. Proteome Res. 12:547–558.

46. Martin, A., T.A. Baker, and R.T. Sauer. 2008. Protein unfolding by a AAA+ protease is dependent on ATP-hydrolysis rates and substrate energy landscapes. Nat. Struct. Mol. Biol. 15:139–145.

47. Peterson, C.N., I. Levchenko, J.D. Rabinowitz, T.A. Baker, and T.J. Silhavy. 2012. RpoS proteolysis is controlled directly by ATP levels in Escherichia coli. Genes Dev. 26:548–553.

48. Nager, A.R., T.A. Baker, and R.T. Sauer. 2011. Stepwise unfolding of a β barrel protein by the AAA+ ClpXP protease. J. Mol. Biol. 413:4–16.

49. Ye, F., J. Zhang, R. Hilgenfeld, D. Li, X. Zhang, H. Jiang, R. Zhang, C.-G. Yang, L. Li, C. Luo, J. Lu, X. Kong, and H. Liu. 2013. Helix Unfolding/Refolding Characterizes the Functional Dynamics of Staphylococcus aureus Clp Protease. J. Biol. Chem. 288:17643–17653.

50. Kimber, M.S., A.Y.H. Yu, M. Borg, E. Leung, H.S. Chan, and W.A. Houry. 2010. Structural and Theoretical Studies Indicate that the Cylindrical Protease ClpP Samples Extended and Compact Conformations. Structure. 18:798–808.

51. Sprangers, R., A. Gribun, P.M. Hwang, W.A. Houry, and L.E. Kay. 2005. Quantitative NMR spectroscopy of supramolecular complexes: Dynamic side pores in ClpP are important for product release. Proc. Natl. Acad. Sci. U. S. A. 102:16678–16683.

52. Zhang, J., F. Ye, L. Lan, H. Jiang, C. Luo, and C.G. Yang. 2011. Structural switching of Staphylococcus aureus Clp protease: A key to understanding protease dynamics. J. Biol. Chem. 286:37590–37601.

53. Kang, S.G., M.R. Maurizi, M. Thompson, T. Mueser, and B. Ahvazi. 2004. Crystallography and mutagenesis point to an essential role for the N-terminus of human mitochondrial ClpP. J. Struct. Biol. 148:338–352.

54. Martín, Á.S., P. Rodriguez-Aliaga, J.A. Molina, A. Martin, C. Bustamante, and M. Baez. 2017. Knots can impair protein degradation by ATP-dependent proteases. Proc. Natl. Acad. Sci. U. S. A. 114:9864–9869.

55. Sriramoju, M.K., Y. Chen, Y.T.C. Lee, and S.T.D. Hsu. 2018. Topologically knotted deubiquitinases exhibit unprecedented mechanostability to withstand the proteolysis by an AAA+ protease. Sci. Rep. 8:1–9.

56. Sivertsson, E.M., S.E. Jackson, and L.S. Itzhaki. 2019. The AAA+ protease ClpXP can easily degrade a 3 1 and a 5 2-knotted protein. Sci. Rep. 9:1–14.

57. Joshi, S.A., G.L. Hersch, T.A. Baker, and R.T. Sauer. 2004. Communication between ClpX and ClpP during substrate processing and degradation. Nat. Struct. Mol. Biol. 11:404–411.

58. Kuo, Y.W., and J. Howard. 2021. Cutting, Amplifying, and Aligning Microtubules with Severing Enzymes. Trends Cell Biol. 31:50–61.

59. Ryu, J.K., R. Jahn, and T.Y. Yoon. 2016. Review: Progresses in understanding N-ethylmaleimide sensitive factor (NSF) mediated disassembly of SNARE complexes. Biopolymers. 105:518–531.

60. Kardon, J.R., J.A. Moroco, J.R. Engen, and T.A. Baker. 2020. Mitochondrial clpx activates an essential biosynthetic enzyme through partial unfolding. Elife. 9:1–20.

61. Puchades, C., C.R. Sandate, and G.C. Lander. 2020. The molecular principles governing the activity and functional diversity of AAA+ proteins. Nat. Rev. Mol. Cell Biol. 21:43–58.

62. Classen, M., S. Breuer, W. Baumeister, R. Guckenberger, and S. Witt. 2011. Force spectroscopy of substrate molecules en route to the proteasome’s active sites. Biophys. J. 100:489–497.

63. Lee, H., B. Pelz, J.M. Ferrer, T. Kim, M.J. Lang, and R.D. Kamm. 2009. Cytoskeletal deformation at high strains and the role of cross-link unfolding or unbinding. Cell. Mol. Bioeng. 2:28–38.

